# Fine-scale resolution and analysis of runs of homozygosity in domestic dogs

**DOI:** 10.1101/315770

**Authors:** Aaron J. Sams, Adam R. Boyko

## Abstract

Inbreeding and consanguinity leave distinct genomic traces, most notably long genomic tracts that are identical by descent and completely homozygous. These runs of homozygosity (ROH) can contribute to inbreeding depression if they contain deleterious variants that are fully or partially recessive. Several lines of evidence have been used to show that long (> 5 megabase (Mb)) ROH are disproportionately likely to harbor deleterious variation, but the extent to which long versus short tracts contribute to autozygosity at loci known to be deleterious and recessive has not been studied.

In domestic dogs, nearly 200 mutations are known to cause recessive diseases, most of which can be efficiently assayed using SNP arrays. By examining genome-wide data from over 200,000 markers, including 150 recessive disease variants, we built high-resolution ROH density maps for nearly 2,500 dogs, recording ROH down to 500 kilobases. We observed over 500 homozygous deleterious recessive genotypes in the panel, 90% of which overlapped with ROH inferred by GERMLINE. Although most of these genotypes were contained in ROH over 5 Mb in length, 14% were contained in short (0.5 - 2.5 Mb) tracts, a significant enrichment compared to the genetic background, suggesting that even short tracts are useful for computing inbreeding metrics like the coefficient of inbreeding estimated from ROH (*F_ROH_*).

In our dataset, *F_ROH_* differed significantly both within and among dog breeds. All breeds harbored some regions of reduced genetic diversity due to drift or selective sweeps, but the degree of inbreeding and the proportion of inbreeding caused by short versus long tracts differed between breeds, reflecting their different population histories. Although only available for a few species, large genome-wide datasets including recessive disease variants hold particular promise not only for disentangling the genetic architecture of inbreeding depression, but also evaluating and improving upon current approaches for detecting ROH.

## Introduction

Chromosomal segments that are homozygous by descent (autozygous) are a hallmark of inbreeding. Close consanguineous matings typically result in offspring with several long runs of homozygosity (ROH), while matings between more distant shared relatives (as often occurs in small or bottlenecked populations) produce a distribution of ROH skewed towards shorter tract lengths (see, for example Figure 2 in (Howrigan et al. 2011)). Most organisms of interest contain a wide array of segregating, often rare, recessive or partially recessive variants that can produce a deleterious phenotype when exposed as homozygous on genomic segments of autozygosity. Therefore, the efficient and accurate identification of ROH is of immense interest in the field of genetics, particularly in conservation biology and plant/animal breeding where avoidance of inbreeding depression is of critical importance.

Estimates of autozygosity can be derived from known pedigrees, where the coefficient of inbreeding (*F*) is estimated as half the coefficient of relatedness (*r*) between the parents of an individual (Wright 1922). However, a pedigree-based estimate of *F* merely measures the mean expected autozygosity of an individual and not the true inbreeding level for an individual, which depends on the actual segregation and transmission of chromosomal segments (Hill and Weir 2011; Keller et al. 2011). Furthermore, in many populations, pedigrees may be inaccurate, incomplete, or missing, leading to incorrect or biased estimates of inbreeding (Cassell et al. 2003).

Genetic marker-based *F* estimates can be more accurate than pedigree-based estimates, but estimates based on only a handful of markers are typically less precise than pedigree-based estimates. For example, early molecular approaches to indirectly estimate *F* from microsatellites involved calculations of multi-locus heterozygosity (MLH), *d^2^*, and internal relatedness (IR) (Coulson et al. 1998; Slate and Pemberton 2002; Amos et al. 2001; Coltman et al. 1998). However, several studies later demonstrated that small microsatellite panels are ineffective at accurately estimating *F* (Slate et al. 2004; Balloux et al. 2004; DeWoody and DeWoody 2005).

Dense genotyping from either whole-genome sequencing or array-based genotyping allows for the detection of ROH and the inference of autozygous segments of the genome. Long ROH are indicative of recent identity-by-descent (IBD) and the sum of these tracts is, in theory, the exact inbreeding level of an individual. However, what constitutes a “long” ROH is unclear (Peripolli et al. 2016) and establishing an optimum length threshold is a challenge (McQuillan et al. 2008).

The two parental chromosomes within a diploid individual can be considered IBD at any point, as IBD is ultimately determined by coancestry back to a coalescent event. Such a definition of IBD is unhelpful, particularly because some evidence exists that longer tracts of homozygosity carry disproportionately more deleterious variation than shorter tracts (Szpiech et al. 2013). Generally, the threshold at which a ROH is considered as an IBD segment has been determined empirically (if not arbitrarily) as a tract length that is long enough to likely have been inherited from a recent common ancestry (Ku et al. 2010).

Estimates of inbreeding are correlated with negative fitness consequences in many populations (Lencz et al. 2007; Nalls et al. 2009; Szpiech et al. 2013; Mészáros et al. 2015; Zhang et al. 2015). Precise estimation of inbreeding is particularly important for understanding inbreeding genetic load (Charlesworth and Willis 2009) and the degree to which this load is a consequence of ROH of various sizes (and thus the thresholds by which biologically relevant IBD tracts should be inferred). Previous studies have used functional predictions rather than known deleterious mutations and have led to different conclusions as to whether short or long ROH harbor more deleterious genetic variation (Szpiech et al. 2013; Zhang et al. 2015).

Domestic dogs are an ideal organism in which to study the phenotypic effects of recent inbreeding. Hundreds of dog breeds, each with their own unique genetic history, form closed populations usually characterized by significant levels of autozygosity owing to founder effects, bottlenecks, popular sires, and artificial selection for conformation or performance. Approximately 200 known Mendelian recessive disease variants have been identified in dogs, the majority of which are potential models for human disease (OMIA) and can be tested efficiently with genotyping arrays. While a few of these variants, like the *SOD1* mutation, which predisposes dogs to degenerative myelopathy (a relatively late-onset disorder), are ancient mutations segregating in dozens of breeds, most of these disease variants are found in only one breed, or at most a few related breeds, suggesting they are relatively recent mutations (Awano et al. 2009; Boyko 2011).

If a method to detect ROH is highly accurate, the vast majority of homozygous recessive disease genotypes should occur in regions classified as ROH (and essentially all homozygous genotypes for diseases caused by recent mutations). Furthermore, the distribution of tract lengths for tracts overlapping these genotypes will enable a direct test of whether longer or shorter ROH tracts disproportionately harbor known recessive disease mutations. By incorporating knowledge of known recessive disease variants along with dense genome-wide data, we can better refine and evaluate methods of ROH detection, and more precisely investigate patterns of ROH across populations and across genomic regions.

Here, we estimate ROH using two popular methods, PLINK (Purcell et al. 2007; Chang et al. 2015) and GERMLINE (Gusev et al. 2009), to examine the association between ROH and known at-risk genotypes (observed cases of homozygous recessive deleterious genotypes) in domestic dogs. We hypothesize that at-risk genotypes will be highly enriched in ROH regions compared to the non-ROH genomic background. This enrichment can be used to evaluate the sensitivity and specificity of ROH-calling methods and can provide a direct test of whether longer ROH tracts are more or less enriched for these recessive disease variants. We additionally characterize the distribution of ROH in 11 common dog breeds as an example of how the distribution of ROH is influenced by the timing and extent of artificial selection in a breed.

## Results

### Distribution of overlaps of ROH with known homozygous at-risk alleles

In total, 670 dogs whose owners consented to participate in research were homozygous for at least one of 29 Mendelian disease alleles assayed by Embark and considered in our analysis. Of those, we measured the genome-wide distribution of ROH in order to compare it to the set of ROH overlapping a homozygous at-risk allele in our sample. We identified the set of ROH overlapping each occurrence of an at-risk genotype and summarized these tracts in four length categories (< 0.5 Mb -- below ROH detection threshold; 0.5 Mb - 2.5 Mb -- short; 2.5 Mb - 5.0 Mb -- medium; > 5Mb -- long).

We analyzed ROH generated from PLINK and GERMLINE using similar parameters for identifying ROH. In short, we considered ROH >= 0.5 Mb as long as they consisted of at least 41 markers. The results are broadly consistent between these two analyses (Table 1), although GERMLINE was modestly more sensitive, in terms of at-risk genotypes overlapping GERMLINE ROH tracts slightly more often than PLINK-generated ROH tracts, so we focus on results from the former here.

**Table 1.** Analysis of runs of homozygosity (ROH) in dogs carrying homozygous recessive deleterious mutations. ROH > 0.5 Mb (detected by our ROH analysis) harbor recessive deleterious alleles at minimum 28.7X more than ROH < 0.5 Mb. *ROH below our detection threshold

|  | % of all ROH |  | % of *SOD1* |  | % of at-risk (no *SOD1*) |  | Relative risk compared to ROH < 0.5 Mb |  |
| --- | --- | --- | --- | --- | --- | --- | --- | --- |
| ROH Length | PLINK | GERMLINE | PLINK | GERMLINE | PLINK | GERMLINE | PLINK | GERMLINE |
| < 0.5 Mb * | 75.9 | 75.1 | 36.4 | 34.3 | 9.9 | 7.8 | 1.0 | 1.0 |
| 0.5 - 2.5 Mb | 4.5 | 4.7 | 19.4 | 18.7 | 14.9 | 14.7 | 25.7 | 29.8 |
| 2.5 - 5.0 Mb | 3.3 | 3.1 | 15.2 | 14.5 | 11.6 | 11.4 | 27.4 | 35.7 |
| > 5.0 Mb | 16.4 | 17.1 | 29.0 | 32.5 | 63.5 | 66.1 | 29.8 | 37.0 |

\*ROH below our detection threshold

For all at-risk cases excluding SOD1, 92 % of at-risk genotypes had an ROH overlapping the at-risk allele. While the longest ROH in our analysis harbor the majority (66%) of deleterious recessive alleles in this panel of at-risk dogs, short ROH nonetheless harbor known recessive disease homozygous genotypes at a rate nearly 30x higher than stretches of DNA that are not considered ROH. Across all tract lengths that we considered, the relative risk of a ROH carrying a deleterious mutation was similar across classes, suggesting that ROH of all lengths may contribute to inbreeding depression in dogs (see Figure 1, Table 1). To ensure that the deviation of the tracts overlapping at-risk genotypes is not a product of random sampling, we resampled the full distribution of homozygosity tracts from at-risk dogs to mimic the sampling of tracts from at-risk dogs (see Figure 1).

**Figure 1.**
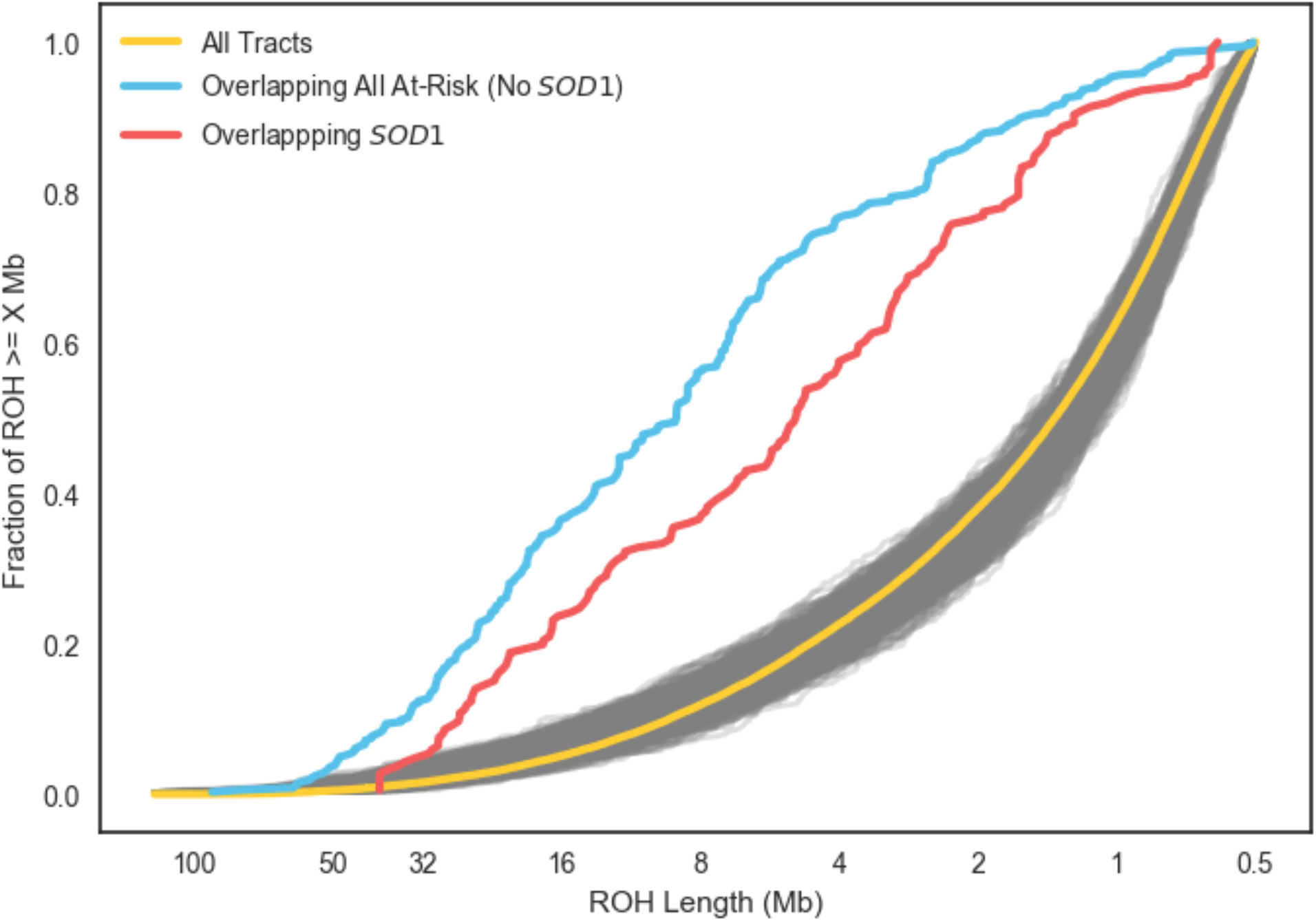
Cumulative density plot of runs of homozygosity (ROH) by length ordered from longest to shortest. Yellow line is all ROH in all dogs at-risk for a deleterious recessive disease, excluding the *SOD1* Degenerative Myelopathy (DM) allele. Blue line is the distribution of ROH overlapping at-risk loci (no *SOD1*) in the same set of dogs. Red is the set of ROH overlapping the *SOD1* DM allele only. Gray lines are 1000 sets of ROH, with the same sample size as the blue line, sampled randomly from the set of all ROH and illustrate that the blue and red sets or ROH are highly non-random samples from the full set of ROH.

Although all the recessive diseases we studied had at-risk genotypes that were highly enriched in ROH tracts, the enrichment was not uniform across these disease variants. Notably for the *SOD1* mutation leading to canine degenerative myelopathy (DM) (Awano et al. 2009), an ancient mutation found in dozens of breeds, we observe a weaker enrichment of at-risk genotypes in long ROH (Figure 1, Table 1).

### Genome-wide distribution of ROH in 11 common dog breeds

We estimated and analyzed the distribution of ROH in 1792 dogs from 11 common dog breeds. First, we calculated *F_ROH_* for all breed dogs and assessed the distribution within breeds for both autosomes and chromosome X. Of the breeds we analyzed Doberman Pinscher had the highest overall levels of *F_ROH_* and Beagle the lowest. In general, *F_ROH_* of chromosome X varied in concert with the autosomes, although some breed (e.g. Doberman Pinscher and Golden Retriever) had somewhat elevated *F_ROH_* on X compare to autosomes while others (e.g. English Bulldog) had lower *F_ROH_* (Figure 2).

**Figure 2.**
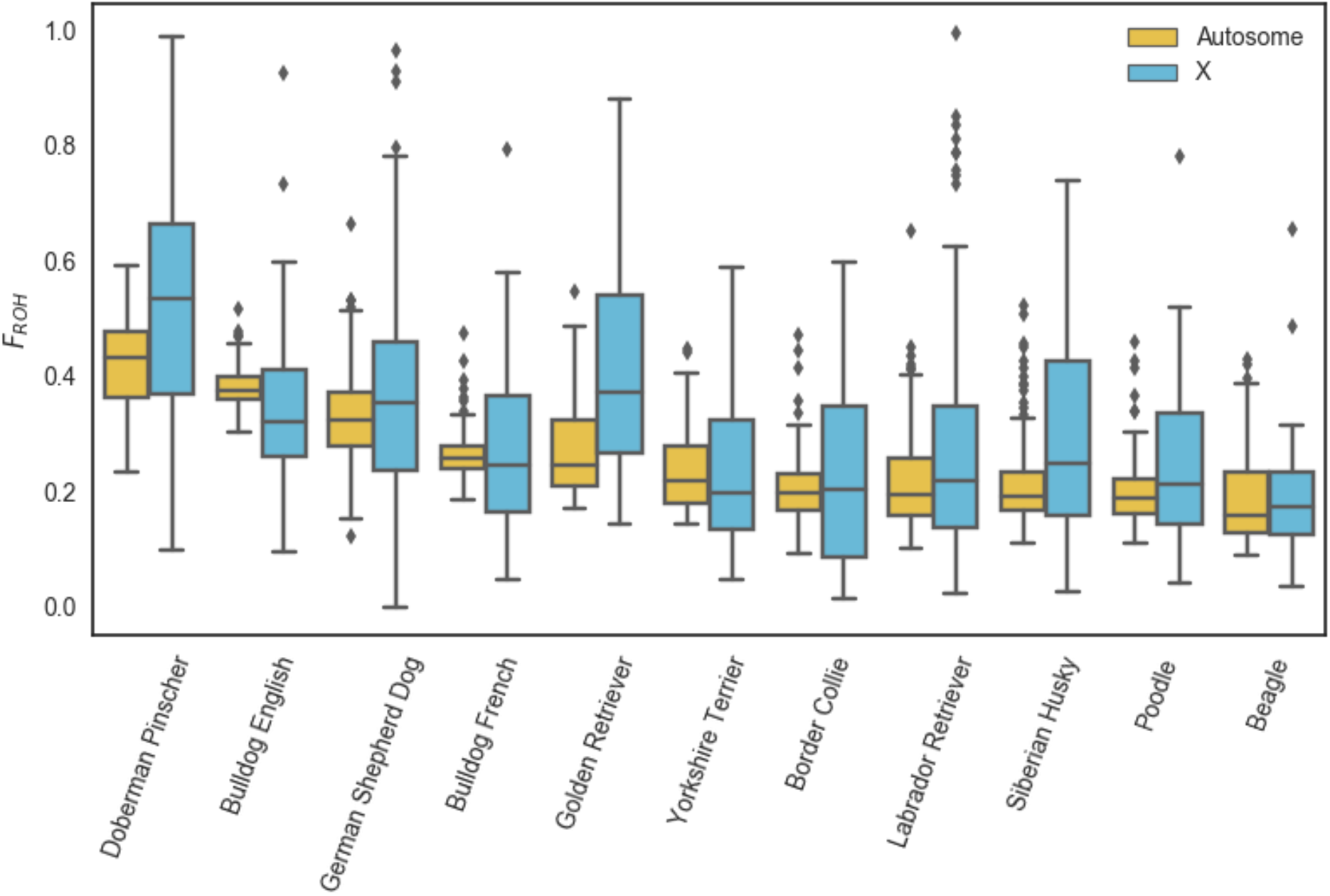
Distribution of *F_ROH_* values across autosomes and the X chromosome for 11 common dog breeds.

For each breed, we also assessed the distribution of ROH by length across all dogs in the breed (Figure 3). These distributions illustrate variation in the timing of diversity loss via inbreeding across breeds. For example, while today Doberman Pinschers have the highest average *F_ROH_* of all breeds in our analysis, the relatively higher fraction of inbreeding in short tracts in English Bulldogs reflects the tremendous bottleneck that occurred in that breed after bull-baiting was banned in the 1830s and the breed was driven to the brink of extinction (Pedersen et al. 2016). Finally, for each breed we calculated a map of the local density of ROH, in other words the fraction of dogs in the breed sample carrying a ROH at each position (Figures S1-11). These maps highlight the deterministic loss of diversity (ROH islands) within breeds, and variation in ROH islands across breeds. Regions of homozygosity associated with fixation of certain variants (e.g. the chr13 *RSPO2* locus in poodles) are clearly evident and are concordant with previously identified homozygosity regions in these breeds (c.f. (Vaysse et al. 2011)). However there is also marked variation in rates of homozygosity outside of these fixed haplotype windows, demonstrating that drift and selection have led to non-uniform diversity loss across the genome in these breeds.

**Figure 3.**
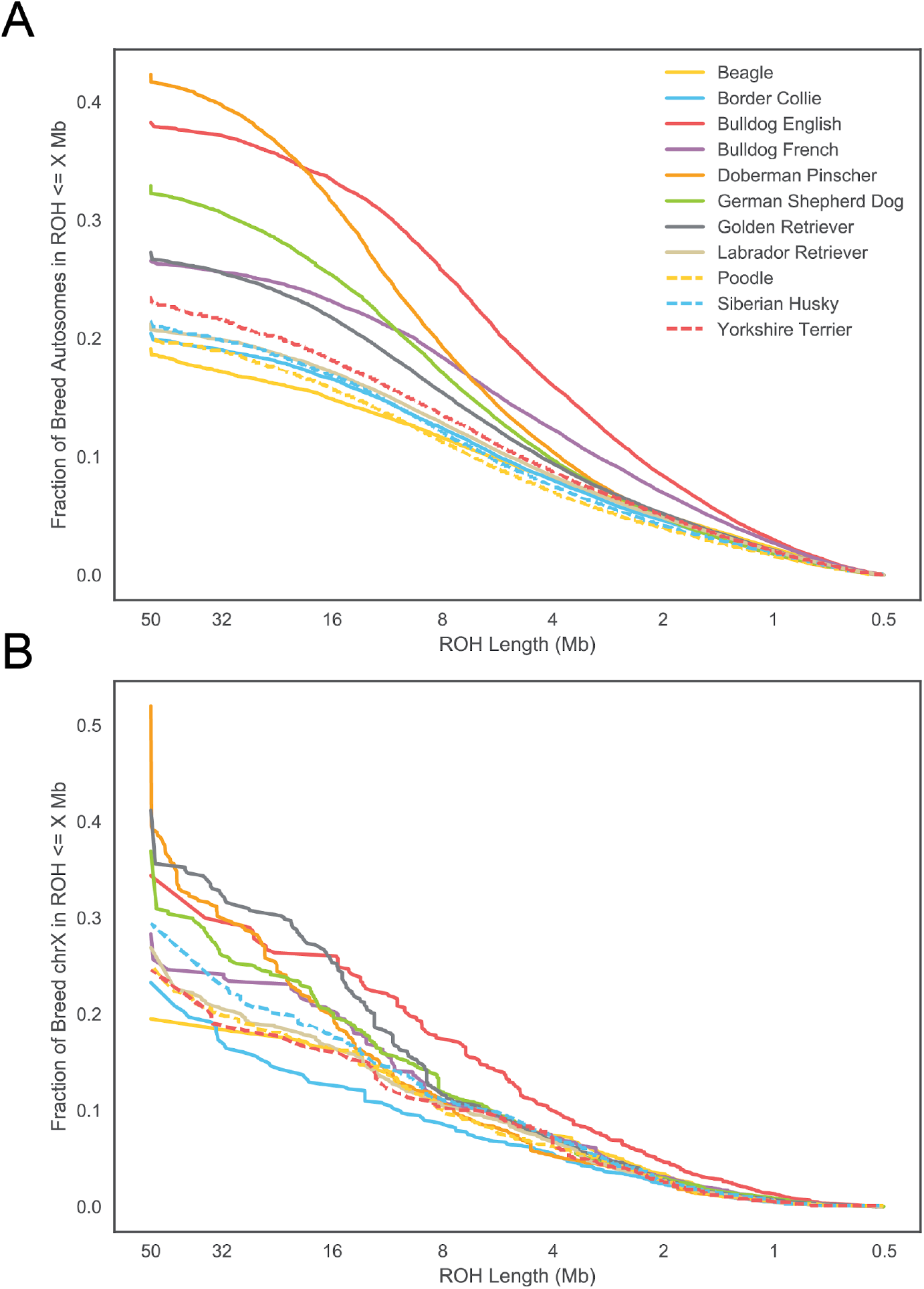
Cumulative density of ROH in 11 common breeds.

## Discussion

Our study is the first to examine patterns of ROH in dogs to a resolution of 500kb. While undoubtedly the sensitivity and specificity of detecting ROH is lower in short (0.5 - 2.5 Mb) tracts compared to long ones, we observe a clear signal of enrichment of known recessive deleterious homozygous genotypes in these regions(Figure 1, Table 1). Thus, these short tracts on aggregate represent a real signal of autozygosity, and furthermore these short tracts, often overlooked in studies of ROH, may be important and measurable contributors to inbreeding depression in dogs and other species. Indeed, 4.8-10.7% of the genomes of these 11 common breeds were covered by ROH tracts between 0.5 to 2.5 Mb (Figure 3A) and these tracts contained 18.7% of the SOD1 and 14.7% of the non-SOD1 known deleterious recessive homozygous genotype calls. Only 34.3% and 7.8% of the SOD1 and non-SOD1 homozygous genotype calls were not detected in ROHs, respectively (Table 1). Because these false negative calls were usually flanked by many homozygous markers (but for less than 500kb) rather than heterozygous markers, we believe they are due to ROH below our detection threshold and not due to point mutations arising on different haplotype backgrounds.

Knowing the extent to which long versus short ROH contribute to inbreeding depression would enable an assessment of the risk posed by recent consanguineous matings, which are often avoidable in dog breeding, and the risk from matings between more distant relatives which is generally unavoidable for purebred dogs. If inbreeding depression in dogs is mainly caused by rare, large-effect recessive (or partially recessive) variants, the contribution of long versus short tracts to inbreeding depression is likely well approximated by the contribution of long versus short tracts to recessive homozygotes at known Mendelian disease alleles, most of which are rare on aggregate, although possibly common in the breed(s) they affect. In contrast if inbreeding depression is mainly caused by common, small-effect recessive or partially recessive variations, the contribution of short tracts to inbreeding depression will be much greater, possibly even greater than the contribution of short tracts to DM risk at the SOD1 locus.

We do see substantial variability in levels of autozygosity between breeds as well as across the genome within a breed. These differences represent the unique history of each breed, and the effect of drift and selection for particular traits over time. At one extreme, many breeds have complete autozygosity in certain windows of the genome. However, genomic regions of high autozygosity that are not completely fixed are also evident in every breed, and efforts to preserve breed diversity should focus on preserving rare haplotypes in these regions rather than rare markers in randomly selected genomic regions. Where fixed haplotypes harbor deleterious variation, marker-assisted crossings and backcrossings to introduce new diversity at the locus or gene-editing techniques like CRISPR to remove the deleterious variant(s) are required to reverse the otherwise irreversible turn of Muller’s ratchet.

Given the intense interest in developing and comparing methods to detect ROH, it is somewhat surprising that these methods are not typically evaluated for their sensitivity and specificity to detect known deleterious recessive mutations (indeed we are aware of no other genomic studies that have done so). While several studies have examined predicted deleteriousness of variants (Szpiech et al. 2013; Zhang et al. 2015), current methods for predicting deleteriousness are almost certainly less accurate than current methods to detect ROH (and predicting recessiveness is even more fraught), making them a poor way evaluate the sensitivity and specificity of ROH detection methods. Being able to evaluate ROH methods in this way, however, is extremely valuable for comparing ROH detection methods and fine-tuning parameters to optimize accurate ROH inference for a population and genomic dataset of interest. Accurate ROH tract detection is invaluable not only for inferring the coefficient of inbreeding and other genetic parameters of interest, but also for accurate reconstruction of population history and identification of relatives.

The dog is an excellent genomic model for evaluating ROH methods as over 200 Mendelian, mostly recessive, variants are known and the requisite genomic resources (e.g. high-quality reference genome and high-density SNP arrays) are available, although at present only the canine array platform used in this study includes both dense genome-wide coverage and probes to assay most of the Mendelian variants known in dogs. Although humans generally have much lower levels of inbreeding than purebred dogs, many more recessive disease variants are known in humans (OMIM), and even denser arrays (including probes for many Mendelian variants) have been used to investigate over 10 million humans to date, largely on commercial DNA testing platforms (https://isogg.org/wiki/Autosomal_DNA_testing_comparison_chart). Thus, high-powered studies looking at many more at-risk loci and many more at-risk individuals with even denser marker panels could potentially be done to further investigate the sensitivity and specificity of various ROH methods and the contribution of different size ROH tracts to inbreeding risk if privacy concerns could be managed and data access granted to researchers in the field. Until then, researchers are encouraged to use this publicly available canine genetic database and to develop similar databases in other model genetic species to improve both the methodology by which ROH is computed and our insights into the genetic architecture of inbreeding depression in these species.

## Acknowledgements

We thank customers of Embark who’s participation made this work possible. We also thank all of the Embark employees who made this work possible, particularly Ryan Boyko, Matt Barton, Erin Chu, Adam Gardner, Tiffany Ho, Andrea Slavney, and Samuel H. Vohr. We also thank the Embark scientific advisory board (in particular Carlos Bustamante for comments on this manuscript), Cornell University, and the Kevin M. McGovern Family Center for Venture Development in the Life Sciences, for their guidance and encouragement. This study was funded by the participants and by Embark.

## Author Contributions

AJS analyzed the data. AJS and ARB jointly wrote the paper.

## Author Information

Conflict of interest statement: AJS and ARB are employees of Embark Veterinary, a canine DNA testing company which offers inbreeding testing using methods similar to those described in this study. ARB is co-founder and part owner of Embark. Correspondence and requests for materials should be addressed to AJS and ARB.

## Methods

### At-Risk Dog Dataset

We queried Embark’s customer database on April 4^th^, 2018 for all dogs whose owners consented to participate in research that were homozygous (at-risk) for recessive deleterious conditions assayed by Embark’s platform. In total, we identified a total of 678 at-risk cases in 670 dogs (some dogs were at-risk for more than one condition). We separated these at-risk cases into two categories: 1) at-risk for *SOD1*-based degenerative myelopathy (Awano et al. 2009), of which we observed 283 at-risk dogs, and 2) at-risk for all 28 other recessive deleterious conditions assayed by Embark, of which we observed 395 at-risk cases across 393 dogs (Table S1).

### Breed Dog Dataset

We queried Embark’s customer database on January 23rd, 2018 for all customer dogs identified as purebred by Embark from the most common 11 breeds and whose owners consented to participate in research. We then used the ‘--genome’ flag in PLINK v1.9 (Chang et al. 2015) to identify pairs of dogs for which the proportion of IBD (PI_HAT) was greater than 0.45 and used these pairs to remove dogs that were potentially related as parent-offspring or full siblings. In total, our final dataset included 1,792 dogs from 11 breeds (Table S2).

### Genotyping & Quality Control

Customer dogs were genotyped on Embark’s custom high-density genotyping platform containing approximately 220,000 markers including all 173,000 markers found on the Illumina CanineHD platform and probes to detect over 160 Mendelian disease variants. SNP filtering using PLINK 1.9 (Chang et al. 2015) was done to ensure genotype concordance rates above 99.99% and missingness rates below 0.1%. Genotype data was phased against a proprietary reference panel and missing data imputed using Eagle2 (Loh et al. 2016). SNP data was also pruned with PLINK to remove markers in close linkage disequilibrium using “--indep-pairwise 200 100 0.90”. After pruning, 170,728 autosomal and 4,395 chrX markers remained, for an average of one marker per 12.8 kb for autosomes (one marker per 28.2 kb on chromosome X).

### Defining Runs of Homozygosity with PLINK

We generated ROH for at-risk dogs in PLINK using software version 1.9 (Chang et al. 2015) (which uses the algorithm from software version 1.07 (Purcell et al. 2007)).

~~~
--homozyg-window-het 0
--homozyg-snp 41
--homozyg-window-snp 41
--homozyg-window-missing 0
--homozyg-window-threshold 0.05
--homozyg-kb 500
--homozyg-density 5000 (set high to ignore)
--homozyg-gap 1000 (set high to ignore)
~~~

### Defining Runs of Homozygosity with GERMLINE

Initially, we attempted to use GERMLINE’s internal filtering to identify ROH >= 0.5 Mb and consisting of at least 41 markers using the following parameters:

~~~
germline -homoz-only -min_m 0.5 -err_hom 0 -err_het 0 -bits 41 -w_extend
~~~

Where -min_m = 0.5 is in units of Megabase-pairs (all measurement in this study was computed in physical distance).

However, we noticed an issue with the germline software in which using the *-w_extend* flag in conjunction with the *-homoz-only* whereby all tracts are extended beyond the first mismatching marker to the end of the next slice (or beginning of the previous slice).

As an alternative, we used the following command in germline to generate preliminary homozygosity tracts for all dogs in this study:

~~~
germline -homoz-only -min_m 0.5 -err_hom 0 -err_het 0 -bits 1 -w_extend
~~~

This identified all segments of the genome >500 kb with no heterozygous markers. We then merged all such segments separated by <50 kb from a neighboring autozygous segment and subsequently removed all merged segments containing fewer than 41 markers (to avoid spurious inference of ROH in regions with few markers). We found this approach superior to allowing a certain set number of heterozygous markers within an ROH for two reasons: (1) requiring no heterozygous variants for at least 500kb vastly improves the specificity for detecting short ROH (500kb - 4000kb), and (2) allowing one or a number of tightly clustered (<50kb) heterozygous variants between two ROHs improves the sensitivity for detecting long ROHs that would otherwise be broken up by genotyping error or copy-number variation (deletions or duplications, most of which are <50kb, can lead to clustered heterozygous at the markers within the structural variant).

### Defining *F_ROH_*

*F_ROH_* was computed as in previous studies (e.g. (McQuillan et al. 2008)) as:

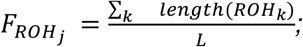

Where ROHk is the kth ROH in individual j’s genome and L is the total length of the genome (or X-chromosome).

